# HaploSight: An alternative approach for Y chromosomal haplogroup identification

**DOI:** 10.1101/2024.03.13.584919

**Authors:** Lukkaet Laoprapaipan, Kittichai Phusawang, Peerapan Laorchairangsi, Arnond Kitnitchee, Hataitip Wachirawanitchkit, Bun Suwanparsert, Pattarapon Promnun

## Abstract

Identification of Y chromosomal haplogroups holds significance in the study of human evolution. In this context, an alternative approach named HaploSight was introduced for identifying Y chromosomal haplogroups. This approach relied on the algorithm of matched and unmatched informative SNPs extracted from the reference phylogenetic tree, implemented in Python. HaploSight demonstrated high accuracy in identifying haplogroups and offered adequate depth in haplogroup analysis. Furthermore, it was compatible with various genotyping arrays, enabling users to work with both sequencing and SNPs dataset.

## Introduction

A phylogenetic tree is a diagram representing evolutionary relationships among various groups of organisms (Yang et al., 2012). Among various DNA markers, the Y chromosome becomes useful for constructing a phylogenetic tree due to its distinctive characteristics (Jobling et al., 2003; Poznik et al., 2013; Wei et al., 2013; Wilson et al., 2014) . As a non-recombining DNA, it provides a clear lineage tracing of ancestry and enables the identification of specific mutations that delineate different branches on the phylogenetic tree. Additionally, as the Y chromosome is paternally inherited, it signifies the most recent common ancestor of all males. Within the Y-chromosomal tree of humans, it is divided into several monophyletic clades called haplogroups. Each haplogroup contains haplotypes identified by a set of unique variants or SNPs. According to previous research, there are currently at least 20 major haplogroups, designed as A-T haplogroups (Hammer et al., 1998; Karafet et al., 2008; Karmin et al., 2015; Poznik et al., 2016; Xue et al., 2009).

Identifying a Y chromosomal haplogroup involves analyzing specific genetic variants on the Y chromosome (Jagadeesan et al., 2021). Several approaches exist for identifying haplogroups based on inputs in formats such as variant calling format and sequencing format (e.g. Jagadeesan et al., 2021; Poznik, 2016; Van et al., 2013). While some specialists are able to develop custom scripts using R or Python in order to align the variants of samples with those in the Y haplogroup tree database, this process might be inconvenient and time-consuming for others. Consequently, there is a need for a tool that efficiently and rapidly pinpoints haplogroups.

The aims of this study were to establish an alternative approach, called “HaploSight”, for processing and accurately identifying the Y haplogroup of men, and to test the performance of the algorithm with different datasets. This tool will be particularly beneficial for researchers who need to rapidly identify Y haplogroups in the large datasets and different types of genotyping arrays.

## Methods

### Algorithm workflow

The first step of our algorithm required a reference phylogenetic tree which could be extracted from either the prior research or generated by the users themselves. Here, we constructed the human whole genome sequence tree from the 1000 Genomes Project, which covers all major haplogroups, under Maximum-Likelihood method. Sequences were downloaded and converted into Phylip format. The program IQtree (Minh et al., 2020) was used to select the best substitution model under Akaike Information Criteria (AIC), and to conduct a phylogenetic tree. The tree was then modified using FigTree (Rambaut, 2018). Following the tree construction, data including tree topology and haplogroups labeled for each node and branch was extracted as a table file. Haplogroups in the tables were annotated using the informative SNPs from various studies to facilitate haplogroup identification. Subsequently, directional assignments were made between paternal haplogroups and their descendants. Each paternal haplogroup was designated as a tree node, with the relationship between a single paternal haplogroup and its descendants represented by the directed arch. These were executed in Python.

The next step involved identifying the haplogroup of samples by matching their SNPs to informative SNPs regarding the annotated table and following the tree directions described previously. For each sample, at each node, the algorithm assessed the number of matched and unmatched SNP alleles at each node. Starting from the bottom nodes of the tree (terminal branches), the algorithm counted the scores for both matched and unmatched SNPs, and then propagated these scores back to the paternal node. Of all descendants, only one route from the paternal node to the child with the highest score was retained. The algorithm iterated until tracing back to the root. The algorithm subsequently selected and presented the route from the root to the terminal branch with the highest score. In case the node containing multiple routes with equal scores, the algorithm stopped and reported from the root to that specific node.

### Performance testing

In order to estimate the accuracy of the HaploSight algorithm, data extracted from 1000 Genomes Project were used as the sample. Initially, the accuracy of the algorithm was evaluated by comparing the haplogroups of whole genome samples predicted by HaploSight to those identified by other algorithms, namely HaploGrouper (Jagadeesan et al., 2021), Y-lineageTracker (Chen et al., 2021), and Y-SNP Haplogroup Hierarchy Finder (Y-Finder; Tseng et al., 2022). For each sample, the deepest nodes identified by these algorithms were compared using density plots. The performance of each algorithm was then tested with the dataset in which the number of SNPs had been reduced. Specifically, the dataset of 1000 Genomes Project was intersected two sets of SNPs in VCF format following different genotyping arrays, which were the Infinium Global Screening Array-24 Kit™ (GSA24) and the Infinium Global Diversity Array-8 Kit™ (GDA8) from Illumina and the Axiom™ Genome-Wide Human Origins (HO) and the Axiom™ Precision Medicine Diversity Array Plus Kit (PMDA) from ThermoFisher Scientific. The haplogroups identified after reducing the number of SNPs were also visualized using density plots. Finally, the distribution of haplogroup change when reducing the number of SNPs of each algorithm was illustrated as a box plot. All analyses were performed using Python.

## Results

The phylogenetic tree of 1000 genome samples was successfully conducted. All haplogroups formed monophyletic. Consequently, this phylogenetic tree served as a reference of our HaploSight algorithm in this study, with haplogroups being extracted and annotated by the informative SNPs for identification purposes. The density plot of deepest haplogroups identified by each algorithm was shown in Figure 1. All algorithms had similar performance in haplogroup identification. Based on our sample, the deepest haplogroups identified by these algorithms reached the 45^th^ level, whereas the highest density of haplogroups was at 30^th^ level. These revealed that the performance of HaploSight was comparable to that of other algorithms.

**Figure 1.**
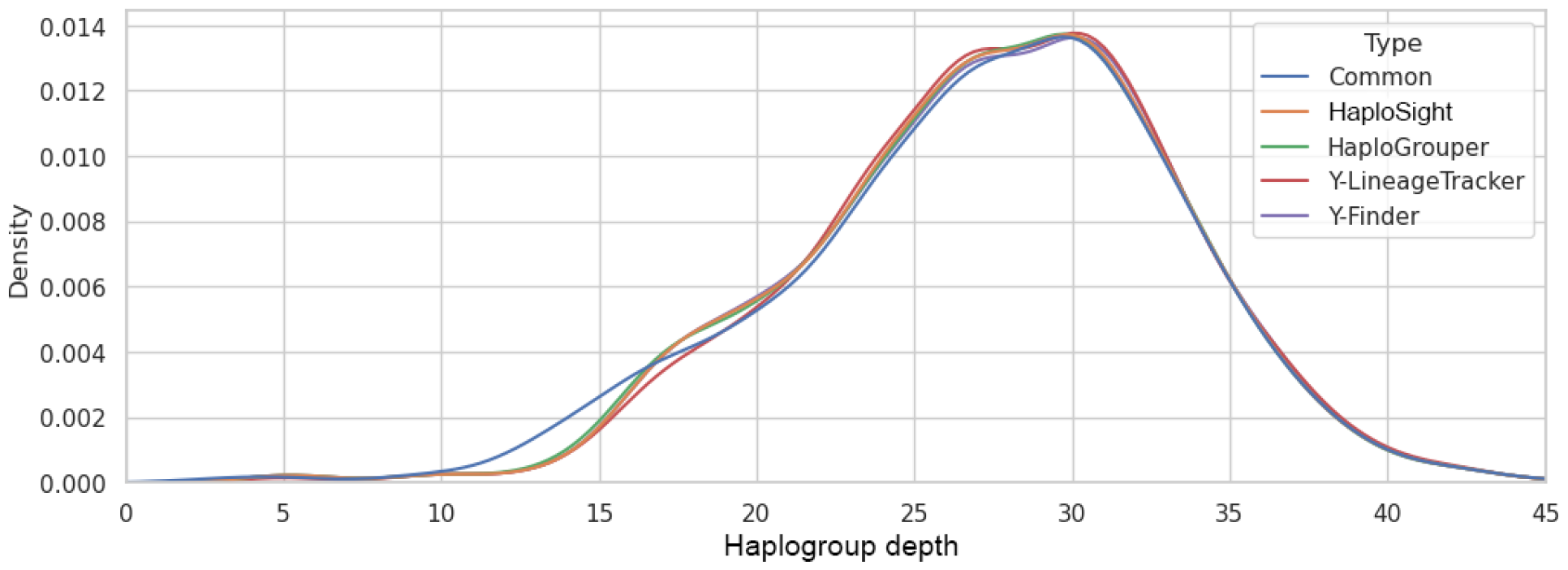
The density plot of haplogroups identified by each algorithm, which were HaploSight, HaploGrouper, Y-LineageTracker, and Y-Finder. The common line represents the common haplogroups that were shared among all algorithms.

The performance of each algorithm when the whole genome dataset was reduced into SNPs dataset following different genotyping arrays was illustrated in figure 2 (a-d). The Y-Finder was excluded from this and subsequent analyses due to unavailability of source code. In general, the deepest haplogroups identified by each algorithm were slightly decreased, reaching approximately the 40^th^ level. When utilizing GDA8 and GSA24 chips from Illumina, the HaploSight could identify haplogroups in comparable depth to other algorithms (figure 2a-b). However, when utilizing the HO and PMDA chips from ThermoFisher Scientific, the patterns of haplogroup depth identified by the HaploSight were more similar to HaploGrouper than to Y-lineageTracker (figure 2c-d).

**Figure 2.**
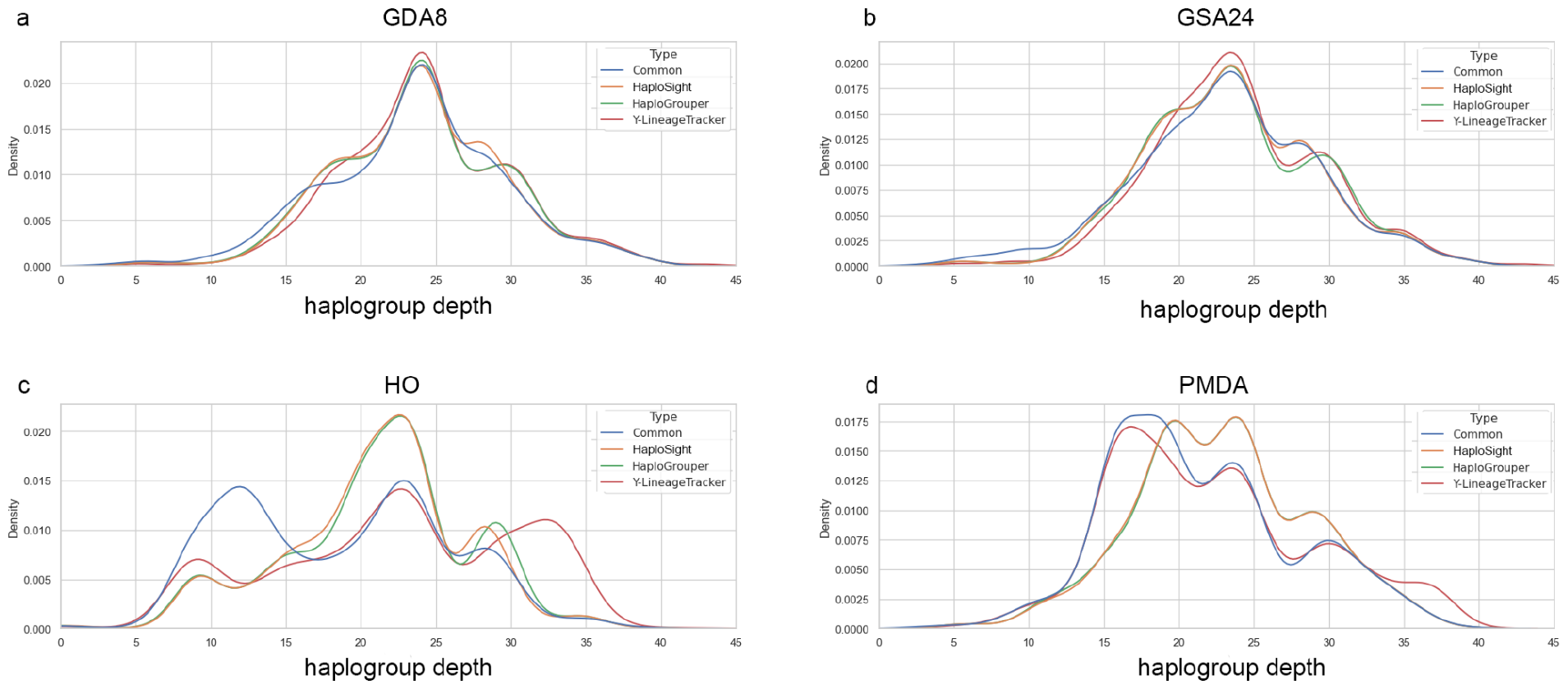
The density plots of haplogroups identified by each algorithms with different genotyping arrays which were a) the Infinium Global Diversity Array-8 Kit™ (GDA8), b) the Infinium Global Screening Array-24 Kit™ (GSA24), c) the Axiom™ Genome-Wide Human Origins (HO), and d) the Axiom™ Precision Medicine Diversity Array Plus Kit (PMDA). The common line represents the common haplogroups that were shared among all algorithms.

The pattern of haplogroup depth identified by the HaploSight using different genotyping arrays was shown in figure 3. Similar to the results observed in figure 1 and 2, the deepest haplogroups identifiable by the HaploSight, as well as the depth with the highest density, were slightly decreased when the dataset was reduced following four genotyping arrays. However, the patterns of haplogroup depth among these four arrays remained similar across these arrays suggesting that the influence of different types of arrays on haplogroup identification by the HaploSight was minimal.

**Figure 3.**
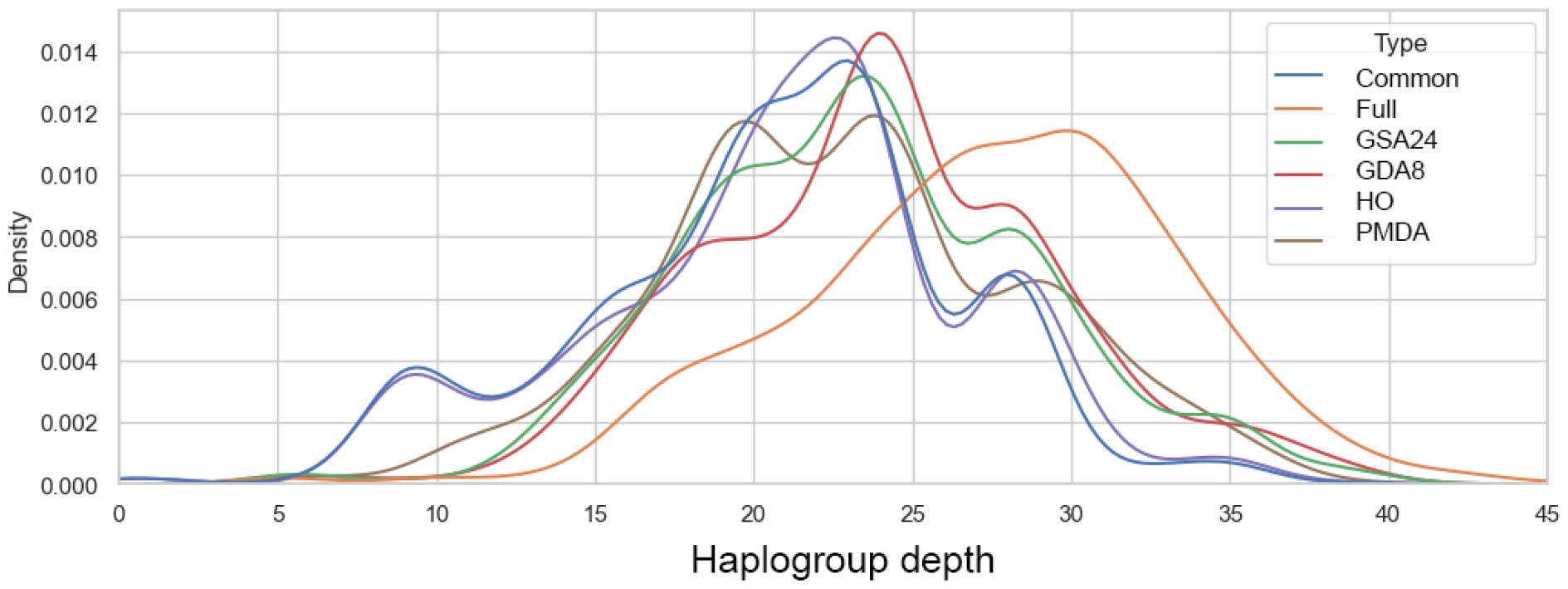
The density plot of haplogroups identified by the HaploSight with different genotyping arrays. The common line represents the common haplogroups that were shared among all arrays.

Finally, the distribution of haplogroup change when the dataset was reduced into four SNPs genotyping arrays was illustrated in figure 4. In this box plot, the x-axis represented the haplogroup depths shared between the whole 1000 genome dataset and each array, while the y-axis illustrated the distribution of haplogroup depth change when employing different arrays. The result indicated that the distribution pattern of HaploSight closely resembled that of HaploGrouper rather than Y-lineageTracker.

**Figure 4.**
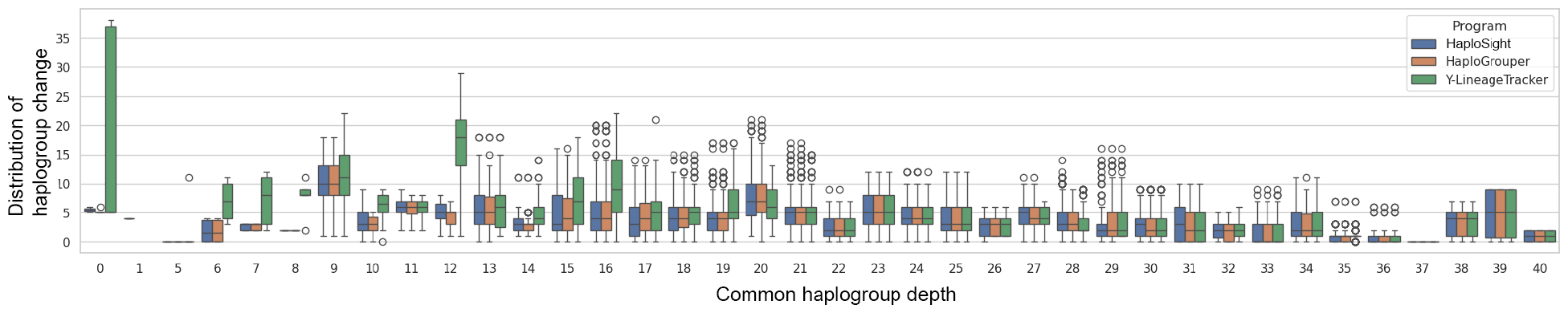
Box plot showing the distribution of haplogroup change when the dataset was reduced into four SNPs genotyping arrays for each algorithm.

Moreover, the change in genotyping arrays had a minimal effect on the haplogroup identification by HaploSight, as evidenced by the low mean and narrow range for each box. This implied that the HaploSight was compatible with various types of genotyping arrays.

## Discussion

Identifying the haplogroup of the human Y chromosome is currently a focal point for scientists due to implication for understanding human evolutionary history (Pinotti et al., 2019; Zeng et al., 2018), and certain haplogroups are also associated with medical genetics concerns (Colaco et al., 2018; Grassman et al., 2019). In this study, we proposed a novel alternative approach for identifying the haplogroup of the Y chromosome called HaploSight.

The HaploSight algorithm possesses several advantages. Firstly, the results obtained from the HaploSight showed successful identification of major haplogroups for all whole genome samples from the 1000 Genomes Projects. The deepest haplogroups of the samples reached depths exceeding 40 levels for both whole genomes and the SNPs dataset. In comparison to the previous studies of the genetic histories of humans, most analyses with other algorithms did not identify haplogroups into such profound levels (e.g. Inoue et al., 2023; Mut et al., 2023; Zieger et al., 2020). According to the Y-chromosomal phylogenetic tree of humans, many haplogroups exhibit further subdivision into a few levels, but only some haplogroups divide further into deep levels such as haplogroup I, J, N, and O (Hammer et al., 1998; Karafet et al., 2008; Karmin et al., 2015; Poznik et al., 2016; Xue et al., 2009). This underscores that the depth identified by the HaploSight was adequate to study human evolution and paternal lineage.

Secondly, the haplogroups identified by the HaploSight showed high accuracy comparable to other algorithms such as HaploGrouper and Y-lineageTracker. The HaploSight algorithm was based on the summation and balancing between matched and unmatched SNPs at each node from the bottom (deepest haplogroups) of the tree upwards. This approach prevented misidentification of haplogroup into incorrect branches and increased the accuracy of identification of major haplogroup. The haplogroups identified by the HaploSight were still accurate even when the DNA markers of samples were reduced from whole genome sequencing into a set of SNPs following two genotyping arrays as indicated by the minimal distribution of haplogroup change (Figure 4). The results supported that the HaploSight was applicable to both FASTA and VCF format and compatible with several and widely used genotyping arrays, such as those from Illumina and ThermoFisher Scientific. Furthermore, HaploSight performed analyses with minimal time-cost, requiring approximately 5 seconds on a 16-core CPU for identifying haplogroups of 1,233 samples (∼ 28 million base pairs of Y-chromosome).

In spite of its strengths, the HaploSight comes with some limitations. Firstly, it depends on the pre-established phylogenetic tree and the informative SNPs that need to be annotated at each haplogroup. In case that the structure of the tree was changed or when applying the algorithm with other trees (rather than the Y-chromosomal tree), it might require more time to construct and annotate the new tree. Moreover, the HaploSight requires a high number of informative SNPs to identify each haplogroup. Consequently, with a lower number of SNPs, the algorithm could likely identify haplogroups into only major haplogroups or into a few depth levels. Reconstruction of phylogenetic trees using more informative SNPs could provide better specificity. Since the phylogenetic tree of the Y chromosome and its markers is still in progress, we intend to update the reference tree on a regular basis. Overall, we anticipate that our algorithm will support users in haplogroup identification.

## Acknowledgements

We would like to thank Jenjit Khudamrongsawat for her comments and editing on our manuscript. We also thank the 1000 Genome Project in providing valuable data for all scientists.

## References

Chen H, et al. 2021. Y-LineageTracker: a high-throughput analysis framework for Y-chromosomal next-generation sequencing data. BMC Bioinformatics. 22, 114.

Colaco S, et al. 2018. Genetics of the human Y chromosome and its association with male infertility. Reprod Biol Endocrinol. 16, 14.

Grassmann F, et al. 2019. International age-related macular degeneration genomics consortium (IAMDGC), Y chromosome mosaicism is associated with age-related macular degeneration. Eur J Hum Genet. 27, 36–41.

Inoue M, et al. 2023. An update and frequency distribution of Y chromosome haplogroups in modern Japanese males. J Hum Genet. Online publication.

Jobling MA, et al. 2003. The human Y chromosome: an evolutionary marker comes of age. Nat. Rev. Genet. 4:598–612.

Hammer MF, et al. 1998. Out of Africa and back again: nested cladistic analysis of human Y chromosome variation. Mol. Biol. Evol. 15, 427–441.

Jagadeesan A, et al. 2021. HaploGrouper: a generalized approach to haplogroup classification. Bioinformatics. 37(4), 570–572.

Karafet TM, et al. 2008. New binary polymorphisms reshape and increase resolution of the human Y chromosomal haplogroup tree. Genome Research. 18, 830–838.

Karmin M, et al. 2015. A recent bottleneck of Y chromosome diversity coincides with a global change in culture. Genome Res. 25, 459–466.

Minh BQ, et al. 2020. IQ-TREE 2: New models and efficient methods for phylogenetic inference in the genomic era. Mol. Biol. Evol. 37, 1530–1534.

Mut P, et al. 2023. Insights into the Y chromosome human diversity in Uruguay. Am J Hum Biol. 35, e23963.

Pinotti T, et al. 2019. Sequences reveal a short beringian standstill, rapid expansion, and early population structure of native american founders. Curr Biol. 29, 149–157.

Poznik GD, et al. 2013. Sequencing Y chromosome resolves discrepancy in time to common ancestor of males versus females. Science. 341, 562–565.

Poznik GD, et al. 2016. Punctuated bursts in human male demography inferred from 1,244 worldwide Y-chromosome sequences. Nat Genet. 48(6), 593–599.

Poznik GD. 2016. Identifying Y-chromosome haplogroups in arbitrarily large samples of sequenced or genotyped men. bioRxiv. 088716.

Rambaut A. 2018. FigTree v1.4.4. Available from http://tree.bio.ed.ac.uk/software/figtree/

Tseng B. et al. 2022. Y-SNP Haplogroup Hierarchy Finder: a web tool for Y-SNP haplogroup assignment. J Hum Genet. 67, 487–493.

Van GA, et al. 2013. AMY-tree: an algorithm to use whole genome SNP calling for Y chromosomal phylogenetic applications. BMC Genomics. 14, 101.

Wei W, et al. 2013. A calibrated human Y-chromosomal phylogeny based on resequencing. Genome Res. 23, 388–395.

Wilson SMA, et al. 2014. Natural selection reduced diversity on human Y chromosome. PLoS Genet. 10:e1004064.

Xue Y, et al. 2009. Human Y chromosome base-substitution mutation rate measured by direct sequencing in a deep-rooting pedigree. Curr. Biol. 19, 1453–1457.

Yang Z, et al. 2012. Molecular phylogenetics: principles and practice. Nat. Rev. Genet. 13, 303–314.

Zeng TC, et al. 2018. Cultural hitchhiking and competition between patrilineal kin groups explain the post-Neolithic Y-chromosome bottleneck. Nat Commun. 9, 2077.

Zieger M, et al. 2020. The Y-chromosomal haplotype and haplogroup distribution of modern T Switzerland still reflects the alpine divide as a geographical barrier for human migration. Forensic Sci. Int. Genet. 48, 102345.

